# Mitochondrial background can explain variable costs of immune deployment

**DOI:** 10.1101/2023.10.04.560830

**Authors:** Megan A. M. Kutzer, Beth Cornish, Michael Jamieson, Olga Zawistowska, Katy M. Monteith, Pedro F. Vale

## Abstract

Organismal health and survival depend on the ability to mount an effective immune response against infection. Yet, immune defence may be energy-demanding, resulting in fitness costs if investment in immune function deprives other physiological processes of resources. While evidence of costly immunity resulting in reduced longevity and reproduction is common, the role of energy-producing mitochondria on the magnitude of these costs is unknown. Here we employed *Drosophila melanogaster* cybrid lines, where several mitochondrial genotypes (mitotypes) were introgressed onto a single nuclear genetic background, to explicitly test the role of mitochondrial variation on the costs of immune stimulation. We exposed female flies carrying one of nine distinct mitotypes to either a benign, heat-killed bacterial pathogen (stimulating immune deployment while avoiding pathology), or to a sterile control, and measured lifespan, fecundity, and locomotor activity. We observed mitotype-specific costs of immune stimulation and identified a positive genetic correlation between lifespan and the proportion of time cybrids spent moving while alive. Our results suggests that costs of immunity are highly variable depending on the mitochondrial genome, adding to a growing body of work highlighting the important role of mitochondrial variation in host-pathogen interactions.

## Introduction

Life history theory predicts that up-regulation of the immune system will come at a physiological cost [1–3]. In line with this prediction, the requirement to reallocate finite energy stores to maintain and activate immune function has been shown to give rise to an array of physiological costs across metazoan taxa [2,4,5]. For instance, there is a strong negative relationship between immune function and reproductive success across species [6], while in many species there is evidence that immune deployment can be costly in terms of reduced longevity [3,7–9]. Additionally, immune activation can trigger significant behavioural changes, such as in locomotor activity, that are intimately intertwined with host energy intake and reallocation [10–12]. Even prokaryotic organisms, which lack what is usually considered to be a conventional immune system, have shown that those with strong pathogen defence mechanisms may experience reduced growth rates [13,14]. Therefore, while the type and form of immunity may vary, its associated fitness costs are likely to be common [5].

While the occurrence of life-history costs associated with immune deployment is well known, the physiological and genetic basis of these trade-offs is rarely understood [15]. Mitochondria, the powerhouses of the cell, generate energy in the form of adenosine triphosphate (ATP) through oxidative phosphorylation (OXPHOS). OXPHOS requires the coordinated action of protein subunits encoded by the nuclear (nDNA) and mitochondrial genomes (mtDNA). Variation in the function of the mitochondria may therefore arise through mutations in nDNA or mtDNA, which can affect signalling between the mitochondrial and nuclear genomes, transcription and translation of mitochondrial proteins, and through the interaction between OXPHOS components of both the nDNA and mtDNA [16]. In the absence of infection or immune deployment, variation in mitochondrial function has been associated with decreased lifespan, changes to locomotor activity and reproductive success, as well as developmental time and weight [17–21]. Thus, we may predict variation in mtDNA to generate, or contribute to, heterogeneity in the immune response [16,22,23]. Following this reasoning, polymorphisms that lead to changes in the efficiency of ATP production may also impact the cost of immune deployment in the form of reduced fecundity, lifespan, or other aspects of host physiology. However, the role of mitochondrial variation on the costs associated with immune stimulation remains poorly understood [22].

Here we asked if variation in the mitochondrial genomes in nine cytoplasmic hybrid lines (cybrids) impacted the expression of two life-history traits, survival and fecundity, as well as locomotor activity in immune stimulated female fruit flies (*D. melanogaster*) exposed to heat-killed *Pseudomonas entomophila*, a natural fly pathogen [24]. mtDNA is inherited maternally, so donor mtDNA can be introgressed (*i*.*e*., repeatedly backcrossed) into a specific nuclear background over multiple generations, ideally resulting in flies with less than 0.01% maternal nDNA, 99.99 % paternal nDNA, and 100 % maternal donor mtDNA [16,20]. We used these cybrid lines to focus on naturally occurring variation in the mitochondrial genome, ensuring that any observed effects were not influenced by the interaction between the mitochondrial and the nucleus. We chose to use heat-killed bacteria to disentangle the effects of immune stimulation and pathology on our life-history readouts: longevity, locomotor activity, and long-term fecundity.

## Materials and Methods

### Generation of cybrid fly lines and rearing conditions

The lines used herein were generated using donor mtDNA from previously described cybrid lines in a wildtype Oregon-R nuclear background [17,20]. The donor mtDNA in each of the eight lines was sampled from naturally occurring geographic variants [20]. To obtain cybrid strains with distinct mitochondrial genomes on the same nuclear background, eight wild-type cybrid strain females (ORT, KSA2, WT5A, BS1, BV1, M2, BOG1, PVM) were backcrossed for 16 generations with males in a W^1118^ nuclear background (Vienna Drosophila Resource Centre). Theoretically, this procedure yield flies with over 99.9% paternal nDNA and 100% maternal donor mtDNA, although this percentage may be lower due to processes such as genetic linkage (Salminen & Vale, 2020). The mtDNA of all strains (except BV1 and w^1118^) had been previously sequenced and the known non-synonymous polymorphisms in OXPHOS complex coding genes are summarised in **Table S1**. All fly lines were maintained at 25±1 °C on a 12 h: 12 h light:dark cycle at 60% humidity, on a standard cornmeal-yeast-sugar diet (14% protein; 1:6 protein:carbohydtate) [25,26]. Experimental flies were raised at constant density for one generation. Ten females and five males were placed in a vial for two days at 25°C, after which, the adults were discarded, and the offspring were left to eclose. Mated female offspring were collected at two to three days post eclosion for both experiments.

### Heat-killed bacteria preparation and confirmation of immune deployment

To stimulate the immune response without a pathogenic infection, we pricked flies with a heat-killed bacterial culture. This procedure is commonly employed in *Drosophila* and other invertebrates to achieve an upregulation of antimicrobial peptide expression in the absence of a viable, replicating pathogen [27–30]. One-hundred microliter aliquots of *Pseudomonas entomophila* were grown overnight in 10 mL of LB broth in an orbital shaker at 140 rpm at 30 °C until the culture reached exponential growth (i.e., OD 0.6 – 0.8). The next morning, cultures were spun down at 2500 rpm/ for 15 minutes at 4°C and the pellet was resuspended in PBS. The OD was then adjusted to 0.1 and further diluted to 0.01 (∼2000 live cells/ fly when pricked [31]). We placed 500 μl aliquots of the dilution into a heat block set to 70°C for 30 minutes and then froze the aliquots for later use. We confirmed that the bacteria had been inactivated by streaking the aliquot out onto a LB agar plate and growing it overnight at 30°C after each experimental block. No viable colony-forming units were detected. Further, all tested fly lines exposed to the heat-killed culture showed an upregulation of the AMP *Diptericin* of between 5 to 50-fold compared to those pricked with sterile PBS (**Figure S1**), confirming that our heat-killed treatment successfully induced immune deployment.

### Locomotor activity and lifespan

Using a randomized block design, we lightly anesthetized 2-3 day old female flies (20-31 flies per genotype per treatment for a total of 472 flies) and pricked them in the thorax with a 5mm needle, dipped in an aliquot of heat-killed *P. entomophila*., or in sterile PBS to control for the effect of pricking. The flies were placed in new vials to recover until being placed in the *Drosophila* Activity monitors (DAM5M), as described previously [11,17,32,33]. Eight DAM5M monitors were set up to measure activity and sleep levels in *Drosophila melanogaster*. Prior to starting the experiment, we prepared food tubes using a 5 % sucrose agar solution. One end of each tube was filled with 2 cm of sucrose-agar. When loading the DAM monitors, flies were anaesthetised using CO_2_ and then placed individually into the tubes of each 32-tube activity monitor at random. The DAM monitors were kept in an incubator (25 °C on a 12 h: 12 h light:dark cycle, 60 % relative humidity) placed to minimise possible disturbance and disruption, and each monitor contained either one empty tube or no tube as negative controls. Each monitor was connected to a laptop placed in the incubator running the DAMSystem3 data collection software. Activity counts and lifespan were recorded in 1-minute bins (Pfeiffenberger et al., 2010) for thirteen days, after which we checked individual survival in each tube.

### Fecundity

234 flies, 13 per line/ treatment) were pricked with PBS or heat-killed *P. entomophila* as described above and then placed individually into vials to monitor egg laying. We transferred the females to fresh food every 24 hours and then counted the eggs laid during the previous day. Daily eggs counts proceeded for 14 days.

### Statistics

Statistical analyses were performed in R version 4.2.2 and RStudio 2022.07.01. We tested for significant interactions and main effects using type 3 Wald χ^2^ tests or likelihood ratio tests [34]. We evaluated model fits using model selection criteria [35] and the *simulateResiduals/ testResiduals* functions in the DHARMa package or the *check_model* function in the performance package. The models are described below and in Table 1.

**Table 1:**
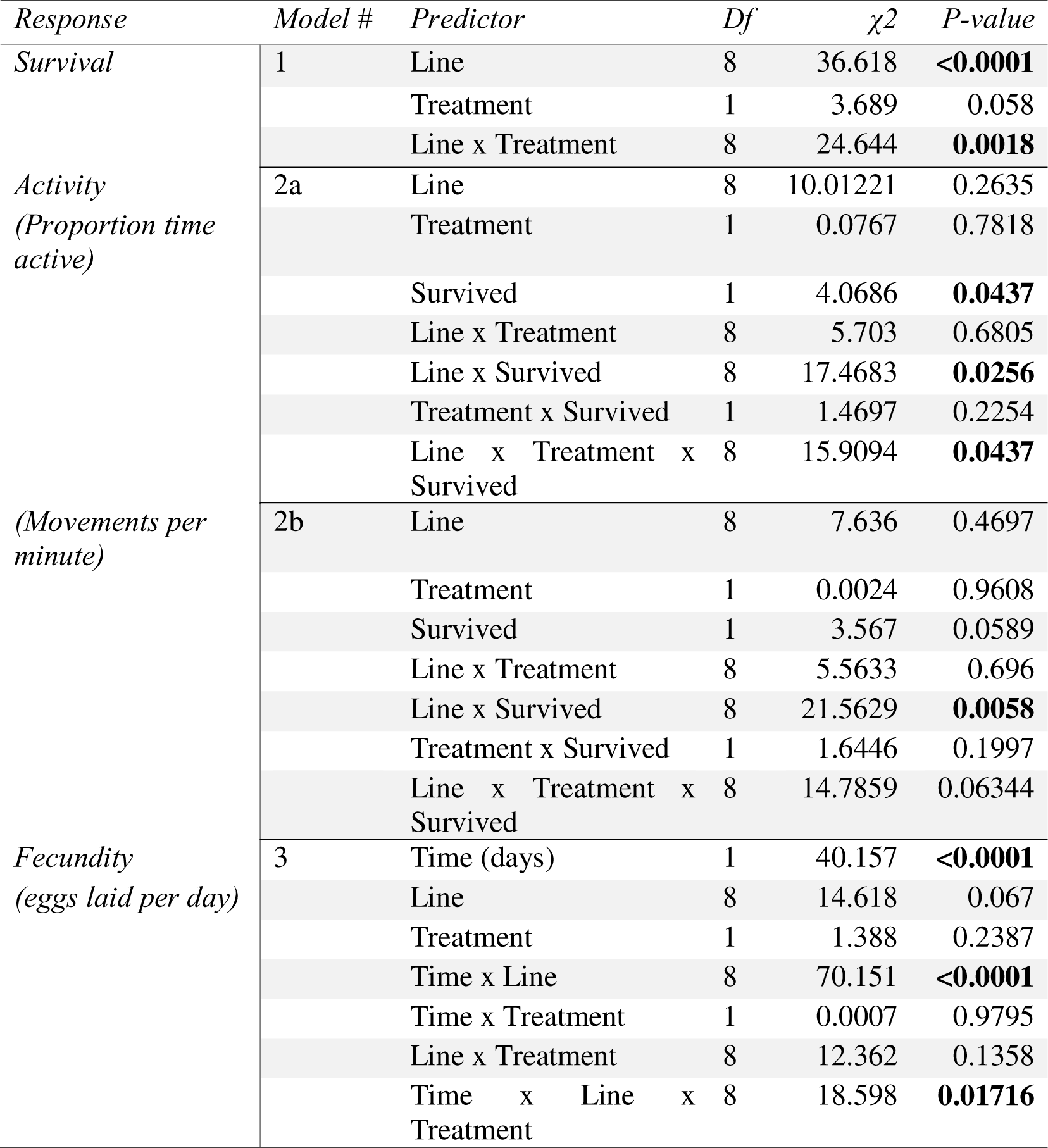
Model outputs.

Survival over the experimental period (Model 1) was analysed using a Cox mixed effects survival model using the *coxme* function in the coxme package [36] with day of death as the response variable and DAM tube nested within monitor and experimental block as random effects. We tested the model without random effects to check whether it fulfilled the assumptions of proportional hazards over time using the *cox*.*zph* function. The model explaining significance variance in survival was:

**Model 1**: *Survival ∼ Line + Treatment + Line × Treatment + (1*|*Block/Monitor/Tube)* We tested for differences in activity levels, measured as the proportion of time spent active (Model 2a) or movements per minute (Model 2b), using linear mixed effects models in the glmmTMB package [37] and tube nested in monitor and experimental block as random effects. We observed that activity levels were largely driven by whether an individual survived, so we included censor/ survived (yes/no) as a factor in our models.

**Model 2a**: *Proportion time active ∼ Line + Treatment + Survived + (Line × Treatment) + (Line × Survived) + (Treatment × Survived) + (Line × Treatment × Survived) + (1*|*Block/Monitor/Tube)*

**Model 2b**: *Movements per minute ∼ Line + Treatment + Survived + (Line × Treatment) + (Line × Survived) + (Treatment × Survived) + (Line × Treatment × Survived) + (1*|*Block/Monitor/Tube)*

We used a generalized linear mixed model with a negative binomial error structure and a quadratic parameterization to evaluate Model 3 using the *glmmTMB* function [37]. We used fecundity, measured as eggs laid per day, as our response variable, time (day) as a covariate, and included FlyID as a random intercept to account for pseudoreplication in the time series.

**Model 3**: *Fecundity ∼ Time + Line + Treatment + (Time × Line) + (Time × Treatment) + (Line × Treatment) + (Time × Line × Treatment) + (1*|*FlyID)*

## Results

### Exposure to heat-killed bacteria induces mitotype-specific mortality rates

We found that individuals pricked with heat killed *P. entomophila* showed increased expression of the AMP Diptericin, confirming several previous studies that this treatment stimulates the immune response despite not containing any viable bacterial cells (see methods for details and Figure S1) [27,29,30]. Flies pricked with heat-killed *P. entomophila* also experienced higher mortality compared to sham-treated flies (0.82 ± 0.13; Figure 1; Model 1), indicating there is a survival cost associated with mounting an immune response. However, the magnitude of the survival cost varied among mitotypes (Figure 1A). Mitochondrial genome differentially affected survival following immune activation when compared to the w^1118^ control (Model 1b, Line x Treatment: p = 0.0018). This effect was largely driven by mtPVM where survival of the sham control was significantly greater than in w^1118^ and by mtBS1, where survival was virtually identical in the sham and heat killed treatments (Figure 1A).

**Figure 1.**
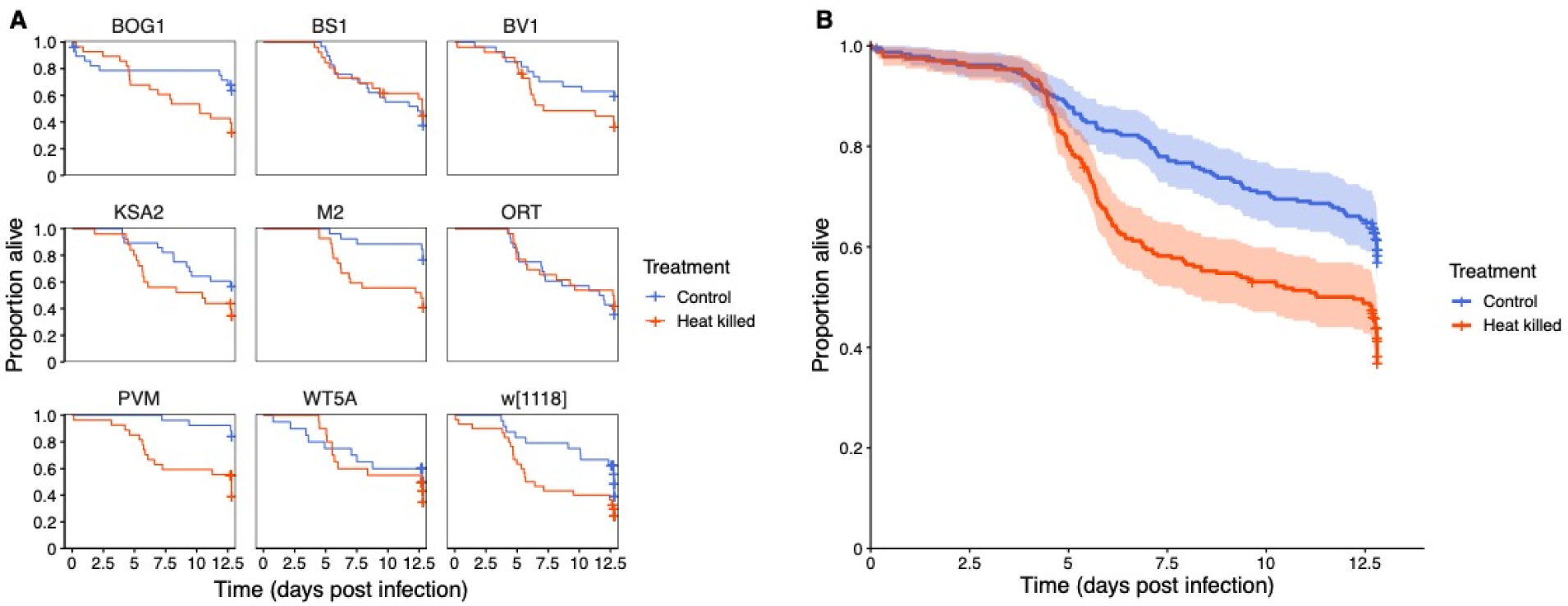
Survival. A) Kaplan-Meier survival curves showing the mortality of flies pricked with either a heat-killed bacterial culture (red) or to a sterile PBS control (blue). Each panel shows a cybrid line with the mitotype indicated above. All lines were introgressed onto a w^1118^ nuclear background for 16 generations. N= 20-31 flies per line per treatment. B) the overall effect of exposure to heat-killed *P. entomophila* on the survival of all flies, averaged across all cybrid backgrounds.

### Fly activity patterns are strong predictors of future survival

Given that different mitotypes are known to impact the extent of fly locomotor activity [17], we tested if costs associated with immune deployment were also apparent in the form of reduced activity and if the magnitude of these varied across mitochondrial lines. We observed that a fly’s survival during the experiment seemed to significantly influence the proportion of time it spent being active while alive (Figure 2A, C). Specifically, we found that individuals that ultimately died tended to spend proportionally less of their lifetime moving while alive, with the magnitude of this decrease depending on mitotype and treatment (Figure 2a; Table 1, Model 2a: Line x Treatment x Survival: p = 0.0437). Additionally, we found a significant and strong positive correlation between the average lifespan of each mitotype and the proportion of time that mitotype remained active while alive (R^2^ = 0.85, p =0.0045, Figure 2e). Surprisingly, flies that died displayed a comparable activity rate (e.g., movements per minute) to flies that survived, regardless of treatment (Figure 2d, Model 2b).

**Figure 2.**
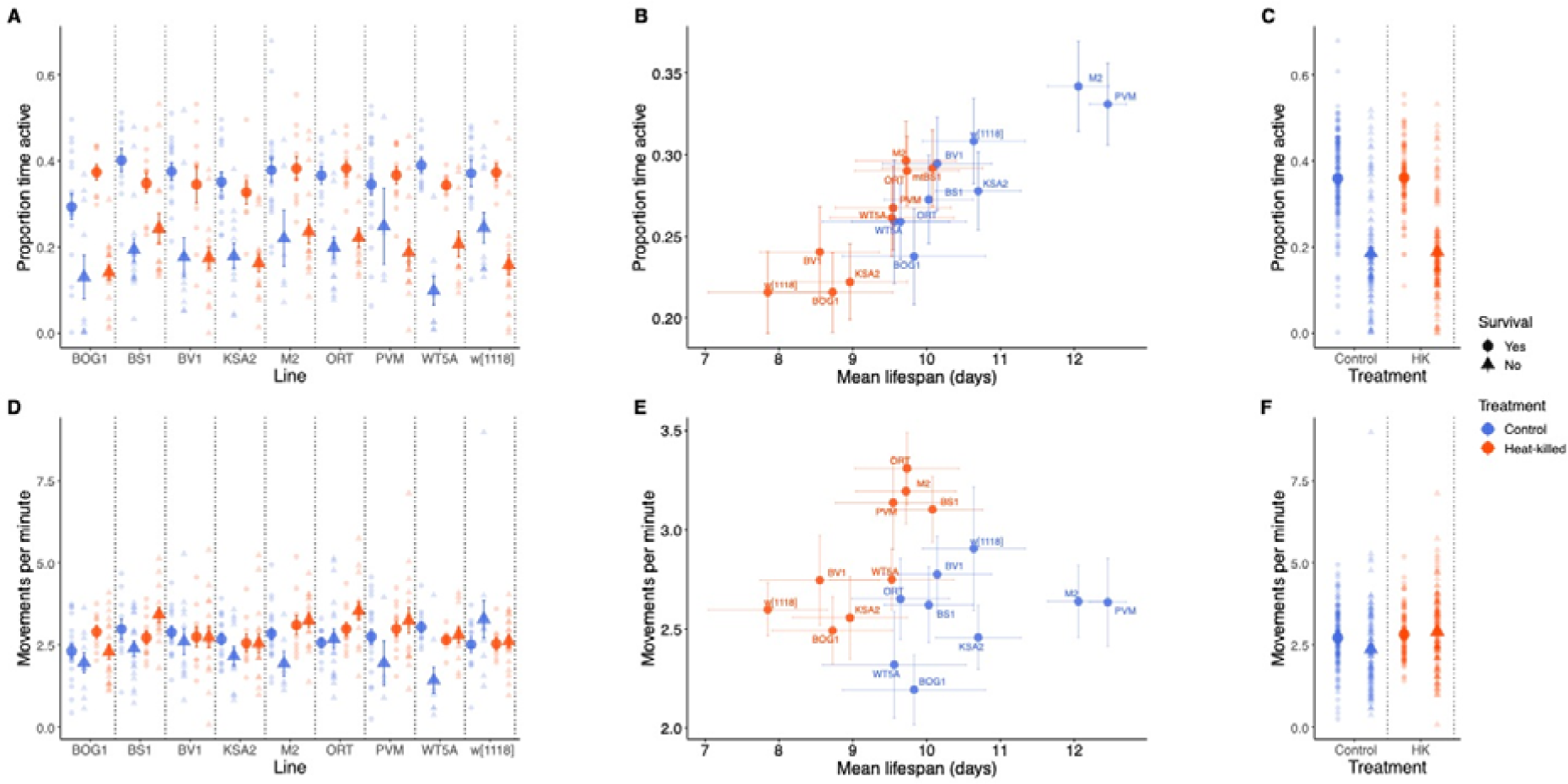
Locomotor activity. **A)** Proportion of time that flies were active over the 13-day experimental period (or their lifetime). Circles represent flies that survived the experimental period and triangles represent flies that died over the thirteen-day period. Each circle or triangle represents a single fly and opaque circles or triangles represent the mean proportion of time spent active +/-1 SE of the mean. Flies inoculated with heat-killed bacteria (HK) are plotted in red-orange and controls are pictured in blue. B) Mean lifespan of each mitotype regressed against the mean proportion of time the mitotype spent active. The heat-killed treatment and controls form distinct clusters. C) The overall effect of exposure to a heat-killed pathogen on the proportion of time individuals were active averaged across all mitotypes. D) Mean movements per minute that flies took over the 13-day experimental period or their lifetime. Circles and triangles represent individual flies that lived or died and means of each mitotype are shown as in (A). E) Mean movements per minute taken by each mitotype regressed against mean lifespan. F) The overall effect of exposure on the mean number of movements averaged across all mitotypes. Colours and symbols are consistent with A-E.

### Egg-laying rate reveals a mitotype-dependent cost of immune stimulation

Costs of immunity are often associated with decreased fecundity in females, so we carried out a separate experiment to test how immune stimulation with heat inactivated bacteria affected fecundity in the cybrid lines over fourteen days. In general, the egg laying rate in flies treated with heat-killed bacteria remained lower than controls from days 1 – 3, stabilizing at day 4 (Figure 3b). However, this time-dependent effect varied according to mtDNA background. Egg laying rate changed over time based on exposure treatment and mitotype (Table 1, Model 3a, Time x Line x Treatment: p = 0.017). All lines exhibited decreased egg laying in the heat-killed treatments one day post exposure, but the magnitude of this effect varied according to mitotype, and the initial cost of immune activation tended to decrease over time. This three-way interaction was driven by BS1, BV1, and WT5A. BS1 and BV1 both recovered fecundity after immune activation compared to w^1118^, whereas fecundity in WT5A remained low in the heat-killed treatment over the two week experimental period. This suggests that mitochondrial background affects immune activation, manifested as reproductive trade-offs.

**Figure 3.**
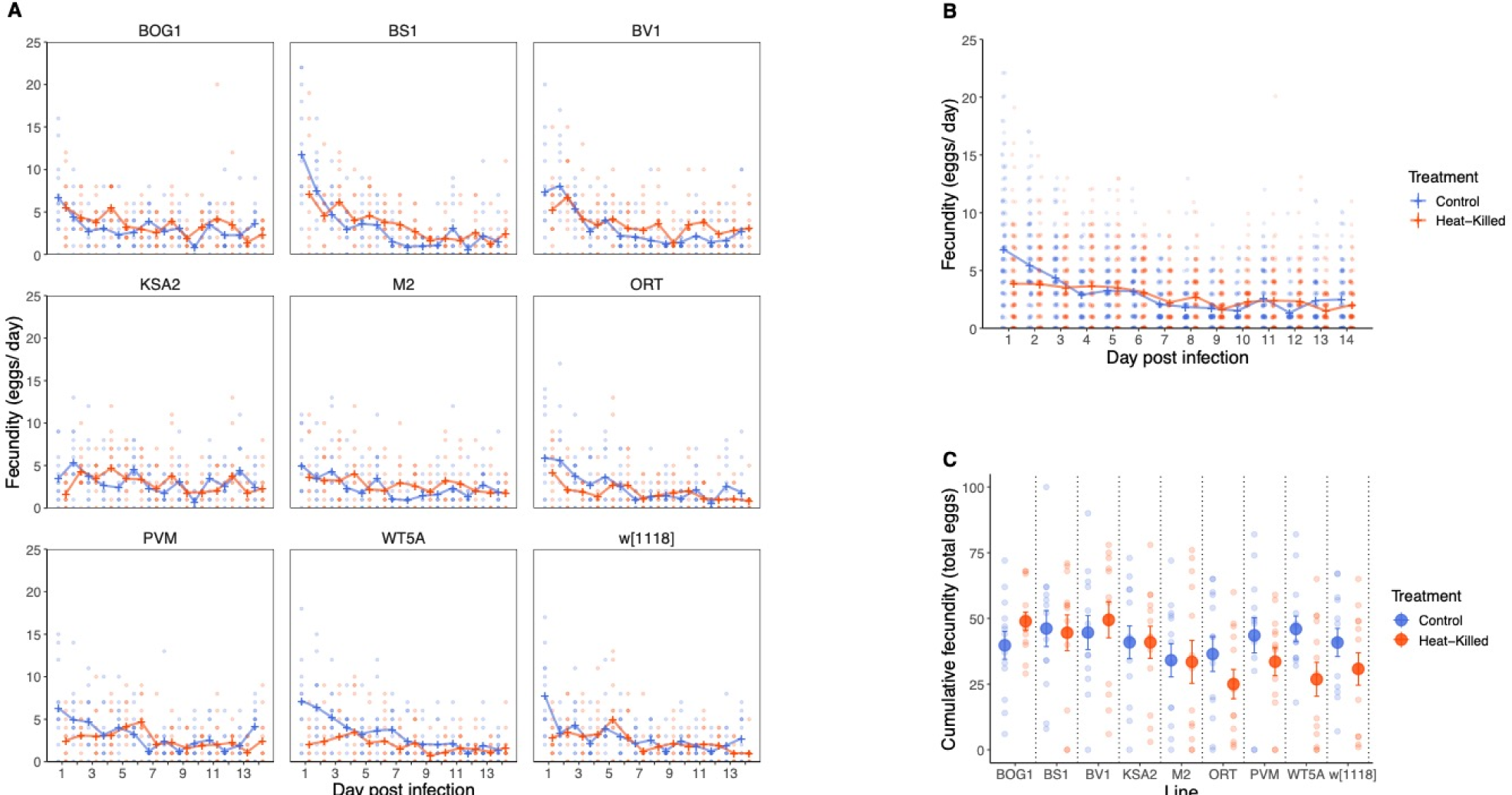
Fecundity differed in response to immune stimulation depending on mitotype over two weeks following challenge. A) Egg laying rate measured as the total eggs laid per day. B) The overall effect of exposure to a heat-killed pathogen on egg laying rate averaged across all mitotypes. C) Cumulative fecundity measured as the total eggs laid per fly over fourteen days. Error bars show 1 SE.

## Discussion

Overall, we found evidence for mitochondrial effects on the cost of immune stimulation in all measured traits. Further, we identified an intriguing positive and strong association between the lifespan of a given cybrid line and the proportion of time it spent moving while alive.

### Mitotype-specific effects on survival

We found cybrid-specific effects on lifespan after immune activation (Figure 1, Table1/ Model 1). Costs of immunity are often associated with reduced longevity in a range of species. For example, immune-stimulation in male field crickets decreased lifespan compared to the controls [7], and a similar result was observed in female eiders [8] and Eurasian collared dove nestlings [9]. A meta-analysis examining tradeoffs between immune activation and life history traits revealed that survival tended to be negatively affected in a range of organisms after immune stimulation by heat-killed bacteria or other antigenic substances [5]. Although much work exists examining the effects of previous exposure to a heat-killed pathogen on survival, many of these studies examine immune stimulation under *ad libitum* laboratory conditions, which may mask life history costs. Costs are often revealed in resource-limited or otherwise challenging environments [2,38,39] and we therefore measured the activity and survival of individual flies within DAM tubes, which is both a resource-limited and stressful environment due to constrained space and reduced access to food. For instance, resource limited bumblebee workers (*Bombus terrestris*) challenged with Lippolysaccharide (LPS), an antigen derived from *E. coli*, or micro-latex beads had significantly reduced lifespans compared to workers with *ad-libitum* access to resources, indicating that life-history costs are sometimes masked by compensatory resource intake [3]. In an Indian meal moth, experimental selection for resistance to a granulosis virus under low or high levels of food resources resulted in reduced resistance in the moths selected on lower resources, suggesting also that immune deployment is costly, and its evolution is resource dependent[38].

### Mitotype-specific effects on fecundity

We measured the mean reproductive output of each cybrid line as the daily number of eggs laid by each fly over a 14-day period, following a single period of mating. We found a significant 3-way interaction between treatment, line, and time (Table 1), revealing a complex effect of immune stimulation on fecundity. Some cybrid lines showed a significant reduction in the number of eggs following stimulation with a heat-killed pathogen, but this effect changed over time and was especially visible within the first 3-5 days of egg-laying (Fig 3). Given that female flies were only mated at the start of the assay, this initial period of egg-laying is therefore the biggest contributor to fitness, as egg-laying rates would be highest during this period and then decrease.

Female fecundity is often – though, not always [25,40,41] -found to be reduced following pathogen exposure, and this reduction is frequently associated with immune-derived trade-offs. In birds, stimulation of the immune response by novel antigen injection in female house martins was accompanied by a decrease in reproductive success compared to noninjected individuals [42]. Studies using female *Drosophila* have also found that mounting an immune response is followed by a decrease in fecundity [2,6] and similarly a post-mating decline in immune function has been observed, presumably due to a reallocation of resources to reproduction [43].

### Immune deployment does not result in changes to activity

In addition to the commonly observed survival and fecundity-based trade-offs, we also measured locomotor activity, a behavioural trait that is likely to be highly energy dependent [17]. We had hypothesized that immune activation would trigger significant behavioural changes because of the relationship between host energy intake and energy reserves. However, activity levels were associated with survival rather than immune status in our study. Conflicting results in other studies suggest that activity levels after immune activation tend to be context dependent. For example, immune stimulation by LPS resulted in a marked decrease in locomotor activity in mice up to 36 hours post inoculation [44] and Eurasian collared dove nestlings injected with LPS had reduced lifespans as a result of predation, which was likely due to reduced activity levels [9]. In contrast, injection with heat-killed *Micrococcus luteus*, a Gram-positive bacterium, induced an increase in activity levels in *D. melanogaster* [12]. Thus, immune activation sometimes comes with significant costs, manifested as behavioural changes, but these tend to be context dependent [45,46] as was evident in the current study.

### Flies that move a lot, live a lot

One intriguing result was the observation of a strong positive genetic correlation between lifespan and the proportion of time flies spent moving (Fig 2B). While we did not set out to test any hypothesis relating the two traits, it became clear during our analysis of locomotor activity that the data generally clustered in two distinct groups within each of the control and heat-killed treatments. Stratifying the data by whether flies had died or remained alive during the experiment showed clearly that these clusters were driven by the much lower proportion of active movement in flies that eventually died (Fig 2). No difference between flies that lived or died was observed in our other measure of activity, movements per minute (Fig 2C,F). The proportion of time spent active should in principle be independent of an individual’s lifespan, so this positive correlation is likely to reflect a true biological effect, rather than any potential statistical artefact of how we measured activity and survival. It is noteworthy that this effect occurs in both heat-killed treated and sham treated flies, so is not clearly a consequence of immune deployment or other immunopathology. Across all tested lines, even within a group of otherwise healthy control flies, a low proportion of time spent active was associated with death later in the experiment (Fig 2A). One potential interpretation is that cybrid lines that are more likely to die are those lines that are already in severe metabolic deficit, and therefore spend a greater proportion of their time inactive. Further work is clearly needed to disentangle the physiological basis of this genetic association. It also suggests that the proportion of time spent active while alive offers a potential useful early warning signal for death [47–49].

### Understanding the mechanisms of mitochondrial effects on life history costs

We have shown evidence that flies carrying distinct mtDNA also experience different levels of costs following immune stimulation, which we measured as reductions in survival, fecundity, or activity. It remains difficult to understand however, which aspect of mitochondrial metabolism contributes to these differences. Mitochondrial function can impact the deployment of immunity in at least three distinct ways [16]: 1) mitochondria play a direct role in immune signalling via intermediates of the mitochondrial tricarboxylic acid (TCA) cycle; 2) mitochondrial metabolism may have a direct antimicrobial action by producing reactive oxygen species (ROS); and 3) mitochondria generate energy by producing ATP during OXPHOS. The non-pathogenic nature of the exposure we employed allows us to disregard the first two sources of variation: all cybrid lines maintained the ability to signal via TCA cycle, and variation in the production of antimicrobial ROS is unlikely to affect the performance of each cybrid line during a benign heat-killed exposure.

This leaves variation in energy production, which we initially predicted would lead to variable costs of immunity. Most cybrid lines we used are known to harbour non-synonymous mutations in several OXPHOS complexes (Table S1, [20]. Mutations in any mitochondrial encoded OXPHOS could affect the total electron transfer chain, potentially causing a decrease in the total production of ATP, with detrimental effects on the expression of life-history traits [20,50]. Changes or loss of mitochondrial function often cause issues with locomotion or flight in *D. melanogaster*. For instance, flies that are mutant for *clueless* and *parkin*, genes associated with altered mitochondrial function, have degenerated flight muscles, display uncoordinated movement, and have shorter lifespans compared to wildtype flies [51].

Recent work on some of the cybrids used in the present work has also found that mitochondrial variation can affect immune deployment directly, by affecting the production of ROS and the proliferation and differentiation of haemocytes during parasitoid wasp infection, particularly in lines with low mitochondrial copy number and non-synonymous substitutions in *cytochrome b* (OXPHOS complex II) and *cytochrome c oxidase subunit 3* (COIII), such as mitotype mtKSA2 [23,50]. In addition, there is some evidence that trade-offs between immunity and fecundity are associated with disruptions in mitochondrial function [22]. Female *D. melanogaster* lacking normal energy metabolism that survived a *Providencia rettgeri* infection had reduced fecundity compared to control genotypes with normal energy metabolism [22]. These findings highlight the importance of mitochondrial function in shaping the life-history costs of immunity.

In summary, we found evidence for costs of immune stimulation measured as fecundity, survival, and activity. The extent of these costs varied according to the mitochondrial genome of each line. Further, we identified an intriguing positive and strong association between the lifespan of a given cybrid line and the proportion of time it spent moving while alive. Our work adds to a growing body of literature highlighting the significant role of mitochondrial variation on the expression of life-history traits [5,18,52,53]. By impacting the costs of immune deployment, mitochondrial variation may therefore also play a role in the maintenance of variation in immunity, with a possibly underappreciated role in host-pathogen evolutionary dynamics [16,54].

## Acknowledgements

We thank Tiina Salminen and Arun Prakash for backcrossing the original mitochondrial genotypes onto the w^1118^ background a year prior to this study. This work was funded by a Leverhulme Trust Research Project grant (RPG-2018-369) awarded to Pedro Vale. The funders had no role in study design, data collection and analysis, decision to publish, or preparation of the manuscript.

## Author contributions

PFV conceived the study, provided funding, reagents and consumables, and designed the experiments together with BC, MJ, OZ and KM. BC, MJ, and OZ carried out experimental work, with assistance and supervision from KMM and PFV. MAMK analysed the data. MAMK and PV wrote the manuscript. All authors read and approved the final version of the manuscript and approved of its submission.

## References

1. Stearns SC. 1992 The Evolution of Life Histories. London: Oxford University Press.

2. McKean KA, Yourth CP, Lazzaro BP, Clark AG. 2008 The evolutionary costs of immunological maintenance and deployment. BMC evolutionary biology 8, 76. (doi:10.1186/1471-2148-8-76)

3. Moret Y, Schmid-Hempel P. 2000 Survival for immunity: the price of immune system activation for bumblebee workers. Science 290, 1166–1168. (doi:10.1126/science.290.5494.1166)

4. McKean KA, Lazzaro BP. 2011 The costs of immunity and the evolution of immunological defense mechanisms. In Mechanisms of Life History Evolution: The Genetics and Physiology of Life History Traits and Trade-Offs (eds T Flatt, A Heyland), pp. 299–310. Oxford: Oxford University Press.

5. Nystrand M, Dowling DK. 2020 Effects of immune challenge on expression of life-history and immune trait expression in sexually reproducing metazoans—a meta-analysis. BMC Biology 18, 135. (doi:10.1186/s12915-020-00856-7)

6. Schwenke RA, Lazzaro BP, Wolfner MF. 2016 Reproduction-immunity trade-offs in insects. Annual Review of Entomology 61, 239–256.

7. Jacot A, Scheuber H, Brinkhof MWG. 2004 COSTS OF AN INDUCED IMMUNE RESPONSE ON SEXUAL DISPLAY AND LONGEVITY IN FIELD CRICKETS. Evolution 58, 2280–2286. (doi:10.1111/j.0014-3820.2004.tb01603.x)

8. Hanssen SA, Hasselquist D, Folstad I, Erikstad KE. 2004 Costs of immunity: immune responsiveness reduces survival in a vertebrate. Proceedings of the Royal Society of London. Series B: Biological Sciences 271, 925–930. (doi:10.1098/rspb.2004.2678)

9. Eraud C, Jacquet A, Faivre B. 2009 SURVIVAL COST OF AN EARLY IMMUNE SOLICITING IN NATURE. Evolution 63, 1036–1043. (doi:10.1111/j.1558-5646.2008.00540.x)

10. Lopes PC, French SS, Woodhams DC, Binning SA. 2021 Sickness behaviors across vertebrate taxa: proximate and ultimate mechanisms. Journal of Experimental Biology 224, jeb225847. (doi:10.1242/jeb.225847)

11. Siva-Jothy JA, Vale PF. 2019 Viral infection causes sex-specific changes in fruit fly social aggregation behaviour. Biology Letters, 630913. (doi:10.1101/630913)

12. Vincent CM, Beckwith EJ, Simoes da Silva CJ, Pearson WH, Kierdorf K, Gilestro GF, Dionne MS. 2022 Infection increases activity via Toll dependent and independent mechanisms in Drosophila melanogaster. PLOS Pathogens 18, e1010826. (doi:10.1371/journal.ppat.1010826)

13. Vale PF, Lafforgue G, Gatchitch F, Gardan R, Moineau S, Gandon S. 2015 Costs of CRISPR-Cas-mediated resistance in Streptococcus thermophilus. Proceedings of the Royal Society B: Biological Sciences 282, 20151270. (doi:10.1098/rspb.2015.1270)

14. Westra ER et al. 2015 Parasite Exposure Drives Selective Evolution of Constitutive versus Inducible Defense. Current Biology 25, 1043–1049. (doi:10.1016/j.cub.2015.01.065)

15. Gupta V, Frank AM, Matolka N, Lazzaro BP. 2022 Inherent constraints on a polyfunctional tissue lead to a reproduction-immunity tradeoff. BMC Biology 20, 127. (doi:10.1186/s12915-022-01328-w)

16. Salminen TS, Vale PF. 2020 Drosophila as a Model System to Investigate the Effects of Mitochondrial Variation on Innate Immunity. Frontiers in Immunology 11. (doi:10.3389/fimmu.2020.00521)

17. Anderson L, Camus MF, Monteith KM, Salminen TS, Vale PF. 2022 Variation in mitochondrial DNA affects locomotor activity and sleep in Drosophila melanogaster. Heredity 129, 225–232. (doi:10.1038/s41437-022-00554-w)

18. Camus MF, Dowling DK. 2018 Mitochondrial genetic effects on reproductive success: signatures of positive intrasexual, but negative intersexual pleiotropy. Proceedings of the Royal Society B: Biological Sciences 285, 20180187. (doi:10.1098/rspb.2018.0187)

19. Clancy DJ. 2008 Variation in mitochondrial genotype has substantial lifespan effects which may be modulated by nuclear background. Aging Cell 7, 795–804. (doi:10.1111/j.1474-9726.2008.00428.x)

20. Salminen TS, Oliveira MT, Cannino G, Lillsunde P, Jacobs HT, Kaguni LS. 2017 Mitochondrial genotype modulates mtDNA copy number and organismal phenotype in Drosophila. Mitochondrion 34, 75–83. (doi:10.1016/j.mito.2017.02.001)

21. Samstag CL, Hoekstra JG, Huang C-H, Chaisson MJ, Youle RJ, Kennedy SR, Pallanck LJ. 2018 Deleterious mitochondrial DNA point mutations are overrepresented in Drosophila expressing a proofreading-defective DNA polymerase γ. PLOS Genetics 14, e1007805. (doi:10.1371/journal.pgen.1007805)

22. Buchanan JL, Meiklejohn CD, Montooth KL. 2018 Mitochondrial Dysfunction and Infection Generate Immunity–Fecundity Tradeoffs in Drosophila. Integrative and Comparative Biology 58, 591–603. (doi:10.1093/icb/icy078)

23. Vesala L, Basikhina Y, Tuomela T, Vale PF, Salminen TS. 2022 Mitochondrial perturbations enhance cell-mediated innate immunity in Drosophila. bioRxiv, 2022.07.19.500574. (doi:10.1101/2022.07.19.500574)

24. Vodovar N, Vinals M, Liehl P, Basset A, Degrouard J, Spellman P, Boccard F, Lemaitre B. 2005 Drosophila host defense after oral infection by an entomopathogenic Pseudomonas species. Proceedings of the National Academy of Sciences of the United States of America 102, 11414–9. (doi:10.1073/pnas.0502240102)

25. Hudson AL, Moatt JP, Vale PF. 2020 Terminal investment strategies following infection are dependent on diet. Journal of Evolutionary Biology 33, 309–317. (doi:10.1111/jeb.13566)

26. Savola E, Montgomery C, Waldron FM, Monteith KM, Vale P, Walling C. 2021 Testing evolutionary explanations for the lifespan benefit of dietary restriction in fruit flies (Drosophila melanogaster). Evolution 75, 450–463. (doi:10.1111/evo.14146)

27. Pham LN, Dionne MS, Shirasu-Hiza M, Schneider DS. 2007 A specific primed immune response in Drosophila is dependent on phagocytes. PLoS Pathogens 3. (doi:10.1371/journal.ppat.0030026)

28. Kutzer MAM, Kurtz J, Armitage SAO. 2019 A multi-faceted approach testing the effects of previous bacterial exposure on resistance and tolerance. Journal of Animal Ecology 88, 566–578. (doi:10.1111/1365-2656.12953)

29. Wen Y, He Z, Xu T, Jiao Y, Liu X, Wang Y-F, Yu X-Q. 2019 Ingestion of killed bacteria activates antimicrobial peptide genes in Drosophila melanogaster and protects flies from septic infection. Developmental & Comparative Immunology 95, 10–18. (doi:10.1016/j.dci.2019.02.001)

30. Prakash A, Fenner F, Shit B, Salminen TS, Monteith KM, Khan I, Vale PF. 2023 The immune regulation and epidemiological consequences of immune priming in Drosophila., 2023.02.22.529244. (doi:10.1101/2023.02.22.529244)

31. Prakash A, Monteith KM, Vale PF. 2023 Negative immune regulation contributes to disease tolerance in Drosophila. bioRxiv (doi:10.1101/2021.09.23.461574)

32. Pfeiffenberger C, Lear BC, Keegan KP, Allada R. 2010 Locomotor Activity Level Monitoring Using the Drosophila Activity Monitoring (DAM) System. Cold Spring Harb Protoc 2010, pdb.prot5518. (doi:10.1101/pdb.prot5518)

33. Vale PF, Jardine MD. 2015 Sex-specific behavioural symptoms of viral gut infection and Wolbachia in Drosophila melanogaster. Journal of Insect Physiology 82, 28–32. (doi:10.1016/j.jinsphys.2015.08.005)

34. Bolker BM, Brooks ME, Clark CJ, Geange SW, Poulsen JR, Stevens MHH, White J-SS. 2009 Generalized linear mixed models: a practical guide for ecology and evolution. Trends in Ecology & Evolution 24, 127–135. (doi:10.1016/j.tree.2008.10.008)

35. Brooks ME, Kristensen K, Darrigo MR, Rubim P, Uriarte M, Bruna E, Bolker BM. 2019 Statistical modeling of patterns in annual reproductive rates. Ecology 100, 1–7. (doi:10.1002/ecy.2706)

36. Therneau TM. 2015 coxme: Mixed Effects Cox Models.

37. Brooks ME, Kristensen K, van Benthem KJ, Magnusson A, Berg CW, Nielsen A, Skaug HJ, Maechler M, Bolker BM. 2017 glmmTMB Balances Speed and Flexibility Among Packages for Zero-inflated Generalized Linear Mixed Modeling. The R journal 9, 378–400.

38. Boots M. 2011 The Evolution of Resistance to a Parasite Is Determined by Resources. The American Naturalist 178, 214–220. (doi:10.1086/660833)

39. Boots M, Begon M. 1993 Trade-Offs with Resistance to a Granulosis Virus in the Indian Meal Moth, Examined by a Laboratory Evolution Experiment. Functional Ecology 7, 528–534.

40. Kutzer MAM, Armitage SAO. 2016 The effect of diet and time after bacterial infection on fecundity, resistance, and tolerance in Drosophila melanogaster. Ecol Evol 6, 4229–4242. (doi:10.1002/ece3.2185)

41. Labbé P, Vale P, Little T. 2010 Successfully resisting a pathogen is rarely costly in Daphnia magna. BMC evolutionary biology 10, 355.

42. Marzal A, Reviriego M, de Lope F, Møller AP. 2007 Fitness costs of an immune response in the house martin (Delichon urbica). Behavioral Ecology and Sociobiology 61, 1573–1580. (doi:10.1007/s00265-007-0389-z)

43. Winterhalter WE, Fedorka KM. 2009 Sex-specific variation in the emphasis, inducibility and timing of the post-mating immune response in Drosophila melanogaster. Proceedings of the Royal Society B: Biological Sciences 276, 1109–1117. (doi:10.1098/rspb.2008.1559)

44. Meneses G et al. 2018 Recovery from an acute systemic and central LPS-inflammation challenge is affected by mouse sex and genetic background. PLOS ONE 13, e0201375. (doi:10.1371/journal.pone.0201375)

45. Lopes PC. 2014 When is it socially acceptable to feel sick? Proceedings of the Royal Society of London B: Biological Sciences 281, 20140218. (doi:10.1098/rspb.2014.0218)

46. Lopes PC, Adelman J, Wingfield JC, Bentley GE, Wingfield JC, Lopes PC, Adelman J. 2012 Social context modulates sickness behavior. Behav Ecol Sociobiol 66, 1421–1428. (doi:10.1007/s00265-012-1397-1)

47. Deb S, Bhandary S, Sinha SK, Jolly MK, Dutta PS. 2022 Identifying critical transitions in complex diseases. J Biosci 47, 25. (doi:10.1007/s12038-022-00258-7)

48. Scheffer M et al. 2009 Early-warning signals for critical transitions. Nature 461, 53–59. (doi:10.1038/nature08227)

49. Trefois C, Antony PM, Goncalves J, Skupin A, Balling R. 2015 Critical transitions in chronic disease: transferring concepts from ecology to systems medicine. Current Opinion in Biotechnology 34, 48–55. (doi:10.1016/j.copbio.2014.11.020)

50. Salminen TS, Cannino G, Oliveira MT, Lillsunde P, Jacobs HT, Kaguni LS. 2019 Lethal Interaction of Nuclear and Mitochondrial Genotypes in Drosophila melanogaster. G3: Genes|Genomes|Genetics 9, 2225–2234. (doi:10.1534/g3.119.400315)

51. Cox RT, Spradling AC. 2009 clueless, a conserved Drosophila gene required for mitochondrial subcellular localization, interacts genetically with parkin. Disease Models & Mechanisms 2, 490–499. (doi:10.1242/dmm.002378)

52. Camus MF, O’Leary M, Reuter M, Lane N. 2020 Impact of mitonuclear interactions on life-history responses to diet. Philos Trans R Soc Lond B Biol Sci 375, 20190416. (doi:10.1098/rstb.2019.0416)

53. Rand DM, Fry A, Sheldahl L. 2006 Nuclear-mitochondrial epistasis and drosophila aging: introgression of Drosophila simulans mtDNA modifies longevity in D. melanogaster nuclear backgrounds. Genetics 172, 329–41. (doi:10.1534/genetics.105.046698)

54. Downey J, Randolph HE, Pernet E, Tran KA, Khader SA, King IL, Barreiro LB, Divangahi M. 2022 Mitochondrial cyclophilin D promotes disease tolerance by licensing NK cell development and IL-22 production against influenza virus. Cell Rep 39, 110974. (doi:10.1016/j.celrep.2022.110974)

