## Supplementary material for "Mitochondrial background can explain variable costs of immune deployment": S1

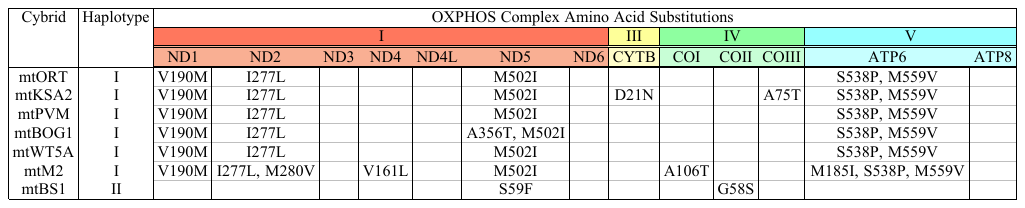

**Table S1.** Deduced amino acid substitutions in the protein coding genes of OXPHOS complexes I, III, IV and V in the *D. melanogaster* mtDNA variants. Table modified from [1].

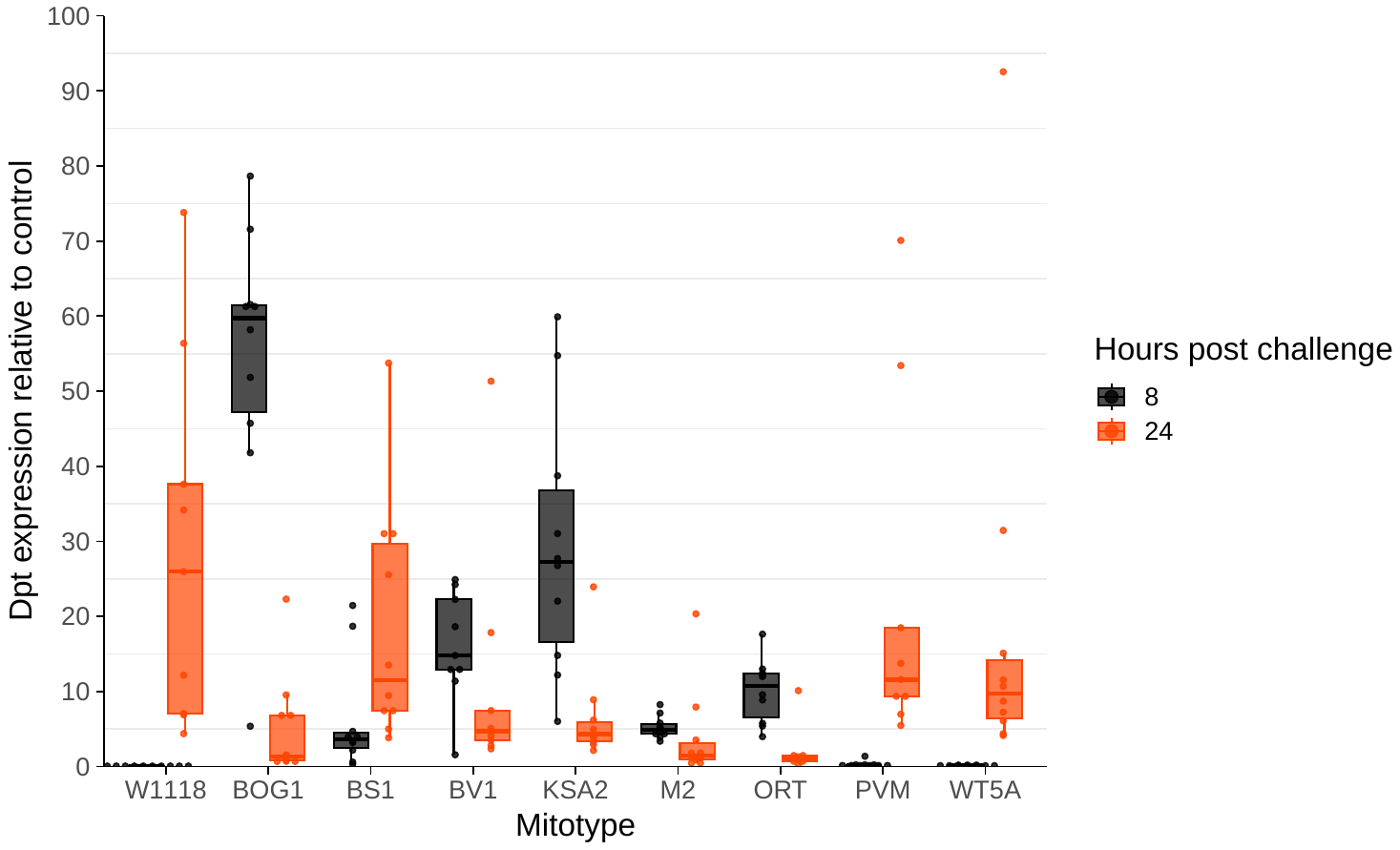

Figure S1. Fold Change in the expression of the antimicrobial peptide Diptericin in flies exposed a heat-killed bacterial pathogen relative to flies exposed to a sterile PBS control. X-axis shows the mitotype of the cybrids tested. Diptericin expression was measured at two time points – 8 (black) and 24 (orange) hours following septic exposure. Primers and qPCR condition are as described in [2].

**Cited references**

1.	Salminen TS, Oliveira MT, Cannino G, Lillsunde P, Jacobs HT, Kaguni LS. 2017 Mitochondrial genotype modulates mtDNA copy number and organismal phenotype in Drosophila. *Mitochondrion* **34**, 75–83. (doi:10.1016/j.mito.2017.02.001)

2.	Prakash A, Fenner F, Shit B, Salminen TS, Monteith KM, Khan I, Vale PF. 2023 The immune regulation and epidemiological consequences of immune priming in Drosophila. BioRxiv, 2023.02.22.529244. (doi:10.1101/2023.02.22.529244)
